# The Specification Game: Rethinking the Evaluation of Drug Response Prediction for Precision Oncology

**DOI:** 10.1101/2024.10.01.616046

**Authors:** Francesco Codicè, Corrado Pancotti, Cesare Rollo, Yves Moreau, Piero Fariselli, Daniele Raimondi

## Abstract

Precision oncology plays a pivotal role in contemporary healthcare, aiming to optimize treatments for each patient based on their unique characteristics. This objective has spurred the emergence of various cancer cell line drug-response datasets, driven by the need to facilitate pre-clinical studies by exploring the impact of multi-omics data on drug response. Despite the proliferation of machine learning models for Drug Response Prediction (DRP), their validation remains critical to reliably assess their usefulness for drug discovery, precision oncology and their actual ability to *generalize* over the immense space of cancer cells and chemical compounds.

This paper shows that the commonly used evaluation strategies for DRP methods learn solutions that optimize an unintended DRP score and fail to predict the proper drug-response activity (”specification gaming”). This problem hinders the advancement of the DRP field, and here we propose a new validation paradigm composed by three Aggregation Strategies (Global, Fixed-Drug, and Fixed-Cell Line) and three train-test Splitting Strategies to ensure a realistic assessment of the prediction performance. We also scrutinize the challenges associated with using IC50 as a prediction label, showing how its close correlation with the drug concentration ranges worsens the risk of misleading performance assessment. We thus propose also an alternative prediction label for DRP which is safer from this perspective.

## 1 Introduction

One of the main goals of precision oncology is to deliver the right drugs in the right doses, on the basis of the specific characteristics of each patient [1]. In order to improve this aspect of the clinical practice we are in need of reliable preclinical models [2]. Large datasets containing drug-response measurements on cancer cell lines have been published, such as the National Cancer Institute 60 (NCI60)[3], the Cancer Cell Line Encyclopedia (CCLE)[4], the Genomics of Drug Sensitivity in Cancer (GDSC)[5], and the Cancer Therapeutics Response Portal (CTRP)[6]. The cell lines in these datasets represent different types of cancer and are usually characterized by various omics data, including sequencing, transcriptomics, proteomics, and methylation data [7].

This data can be used to design computational models that serve as *in silico* alternatives to *in vitro* cell viability screenings [7, 8], providing tailored predictions of drug response across various cell lines. Such Drug Response Prediction (DRP) models would be particularly useful if they could generalize to unobserved drugs or cancer types [9], providing information about potential drug candidates for further analysis, thereby expediting the cancer drug discovery process [9].

Various approaches for DRP in cancer cell lines have been explored, from traditional models like Bayesian Matrix Factorization [10], Random Forests [11], and Support Vector Machines [12] to more recent Neural Networks and Deep Learning techniques [13, 14, 15]. These methods, including Convolutional NNs, Graph NNs and Multimodal DL architectures are capable of handling complex, high-dimensional data and have been used to model drugs, genetic features, and to integrate multiomics data [13, 16, 17, 18, 19, 20, 21, 22].

Aside from the sophistication of the models employed for this task, in this paper we investigate two crucial but often overlooked aspects of DRP, which are 1) the prediction label used to train the models and 2) the validation approach used to evaluate the prediction performance.

For what concerns the first point, most DRP methods focus on the regression of IC50 values to measure drug response [8]. The IC50 value corresponds to the drug concentration necessary to inhibit the viability of 50% of the cells, which is obtained by dose-response curve experiments. These experiments are performed within specific concentration ranges chosen based on the existing knowledge on the target drug [23]. In the paper, we show that the final IC50 values are highly dependent on these ranges, and in particular on the Maximum Concentration (MC) tested. This scenario leads to DRP models that struggle to generalize to new drugs, and they cannot even guess the expected concentration ranges.

The second point relates to the validation approaches used to evaluate the performance of DRP methods. They consist of the combination of how the train and test sets are created (Splitting Strategy), and the way in which the prediction results are aggregated to compute the prediction scores (Aggregation Strategy). Our *in silico* experiments show how the subtleties hidden in the validation of DRP methods can lead to completely misleading performance scores, depending on the characteristics of the datasets used, and how the proper combination of the right Splitting and Aggregation strategies can overcome these issues, by evaluating the model exactly on the kind of task it is designed to solve. For example, since on the most important DRP datasets, including GDSC, CCLE, and CTRP, the type of drug tested is the *main driver* for the variability of the IC50 values, just learning which drugs are generally *strong* or *weak* allows any DRP predictor to *fool* any *global* evaluation metrics based on a simple average of the dose-response values over the entire datasets. Regrettably, this misleading evaluation setting remains prevalent across all current DRP methods. Our study highlights that despite the seemingly impressive global performance metrics, DRP models may still lack any real capability to accurately predict outcomes for novel (previously unseen) cell lines or drugs.

These puzzling results show that, unless stricter evaluation criteria are put in place, specifically targeted for the type of generalization ability that we want to test (drugs or cell lines), the model is able to bypass the conventional evaluation metrics, similarly to what has been shown in other contexts, such as image recognition, cancer driver prediction and Reinforcement Learning [24, 25, 26]. This leads to a *specification gaming* [26] situation, in which the model satisfies the evaluation criteria without attaining the desired outcome. The high performance scores are reached instead by exploiting an unfortunate combination of a peculiar data structure and evaluation metrics that are too generic to be robust to loopholes in the data [25, 26].

To prevent the model from obtaining high-performance scores by just *gaming* the validation specifications of DRP, in this paper we propose different ways to aggregate the predictions (Aggregation Strategies) in order to compute more meaningful evaluation metrics. Each of them is specifically meant to measure a particular generalization ability (towards drugs or cell lines). We also show how the choice of Aggregation Strategy critically depends on how the data are split into training and test subsets (Split Strategy).

The novel validation protocols for DRP that we propose uses AUDRC instead of IC50 and can be specifically targeted to the type of generalization expected from the model under scrutiny (prediction of novel drugs or novel cell lines).

## 2 Results

### 2.0.1 The train-test splitting strategy must simulate realistic prediction scenarios

The observed performances of ML methods depend on the way in which the samples are allocated between training and test sets within the validation procedure of choice, since this defines the underlying assumptions that are tested [8, 27, 28, 25]. To investigate which aspects of the cancer DRP are more challenging, and what level of performances can be realistically expected in real-life settings, in the rest of the paper we analyze the DRP validation problem. Here we start by listing the increasingly stringent strategies available to define the training and test sets:

1. *Random Splits:* This approach is also called Mixed-Set in [8, 29], and it is generally the least challenging, leading to the highest observed performance scores. In this scenario, a randomly selected subset of drug-cell line pairs is excluded from the training set and used as the test set. This train-test Splitting Strategy quantifies how accurate a model is in *filling the gaps* in a drug-cell lines matrix containing some unobserved values. Practically, this would correspond to filling a non-exhaustive screening on a panel of otherwise known cell lines and drugs. In this scenario, the model is *not* evaluated in terms of its ability to generalize to cell lines or drugs for which we completely lack drug-response measurements.
2. *Unseen Cell Lines:* In this case, the train and test splits are made by ensuring that the cell lines in the training set are not present in the test. The test set is constructed by randomly selecting a subset of *cell lines* and *all of* their IC50 values from the whole dataset. To achieve high performance scores in this validation, the models need to be able to generalize to unseen cell lines. With respect to the Random Splits, this therefore increases the difficulty of the prediction task.
3. *Unseen Drugs:* The train and test splits are made ensuring that the drugs that appear in the test set are not present in the training set. To perform well in this setting, the model must be able to generalize well to completely unseen drugs.
4. *Unseen Cell Line-Drug pairs:* This is the most stringent validation setting. In this case, the train and test splits are built to ensure that each of the cell lines and drugs present in the test set are both absent from the training-set. This setting therefore evaluates the ability of the model to generalize at the same time to unseen drugs and cell lines, which should be the ultimate goal of the cancer drug sensitivity prediction field.

These different Splitting Strategies are generally known in DRP literature [9, 8, 30]; however, there is noticeable variability in their actual use [8]. Fig. 3A visually illustrates the generalization difficulty that the DRP models need to overcome, when they are assessed using these strategies.

**Figure 1:**
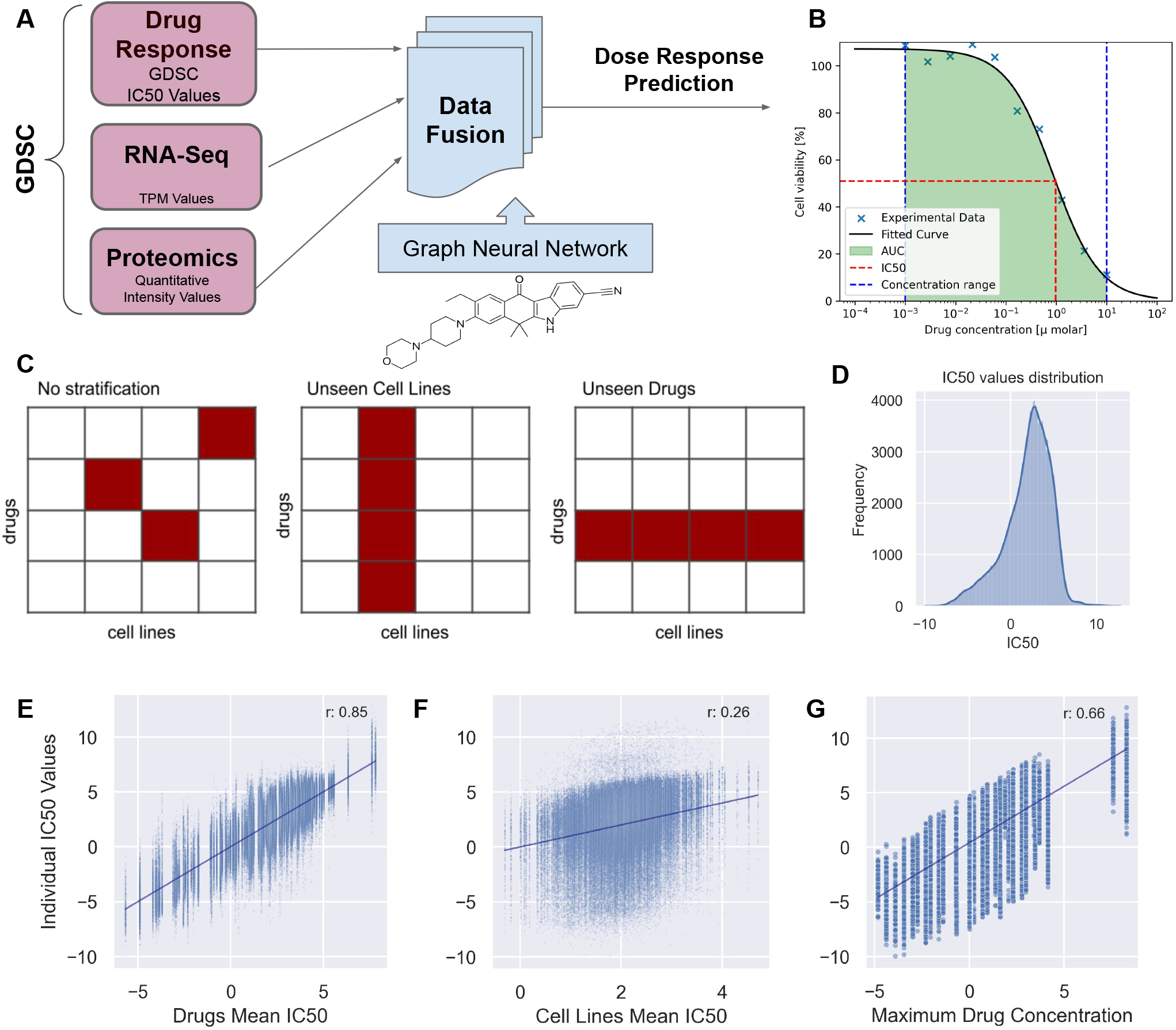
A) *NxtDRP* complete pipeline. B) IC50 values distribution. C) Validation settings: Random Splits, Unseen Cell Lines and Unseen Drugs. Red cells represent the test set. D) IC50 values distribution. E) Individual IC50 values with respect to their relative drug’s mean IC50. F) Individual IC50 values with respect to their relative cell line’s mean IC50. G) Individual IC50 values with respect to the relative drug’s maximum concentration tested for that cell-viabilty experiment. Note: All the IC50 values are relative to the GDSC dataset and are expressed in logarithmic scale.

**Figure 2:**
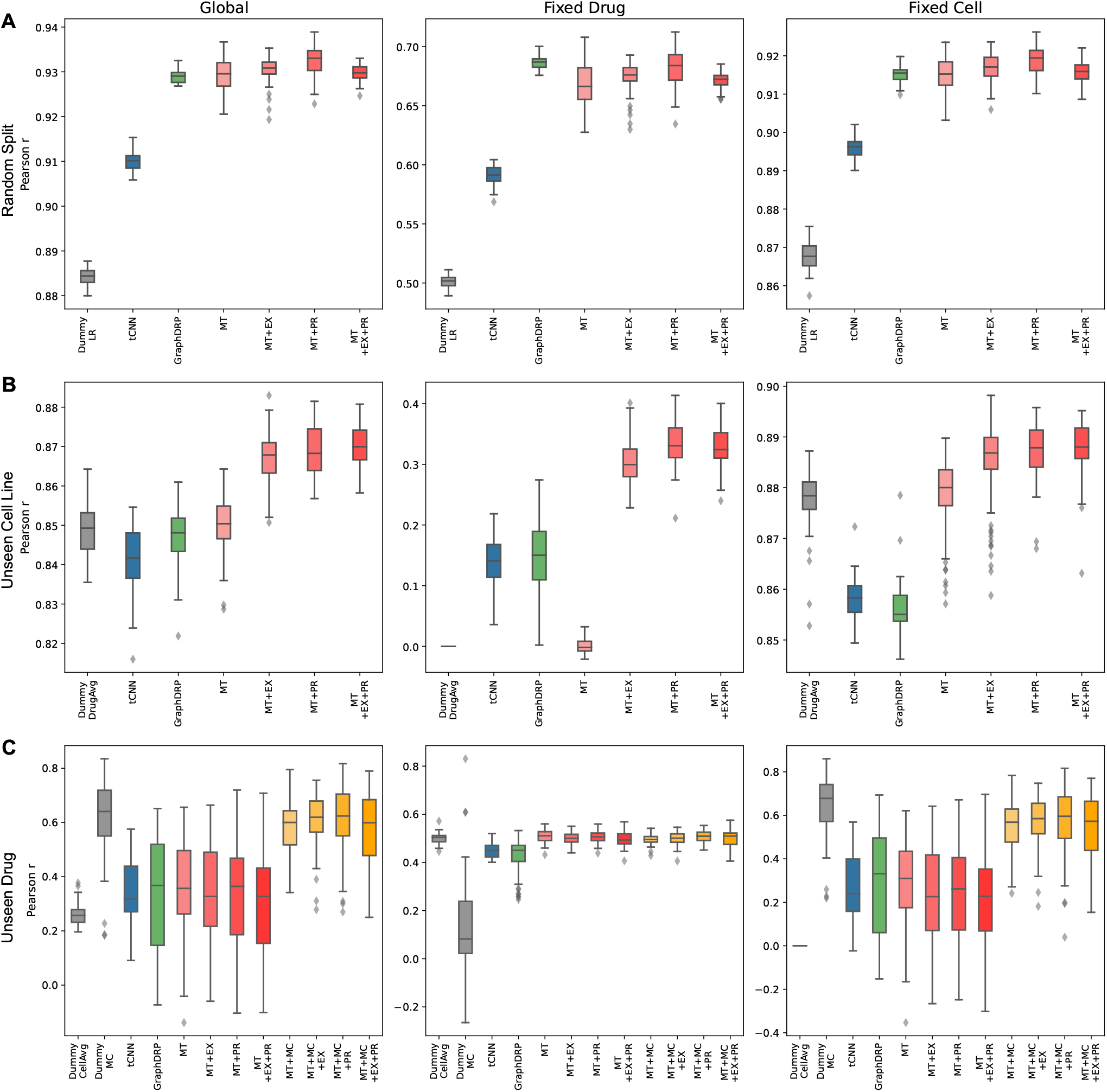
Boxplots showing the distribution of the prediction performances, measured by Pearson’s *r* values, for the tCNN [15], GraphDRP [14], and the NxtDRP models on the GDSC dataset across three Aggregation Strategies (columns) and three Splitting Strategies (rows). The NxtDRP variants MT, PR, and EX denote the omics data utilized: none, Proteomics, and Transcriptomics, respectively (for further details, refer to Fig. 6 B-E).

**Figure 3:**
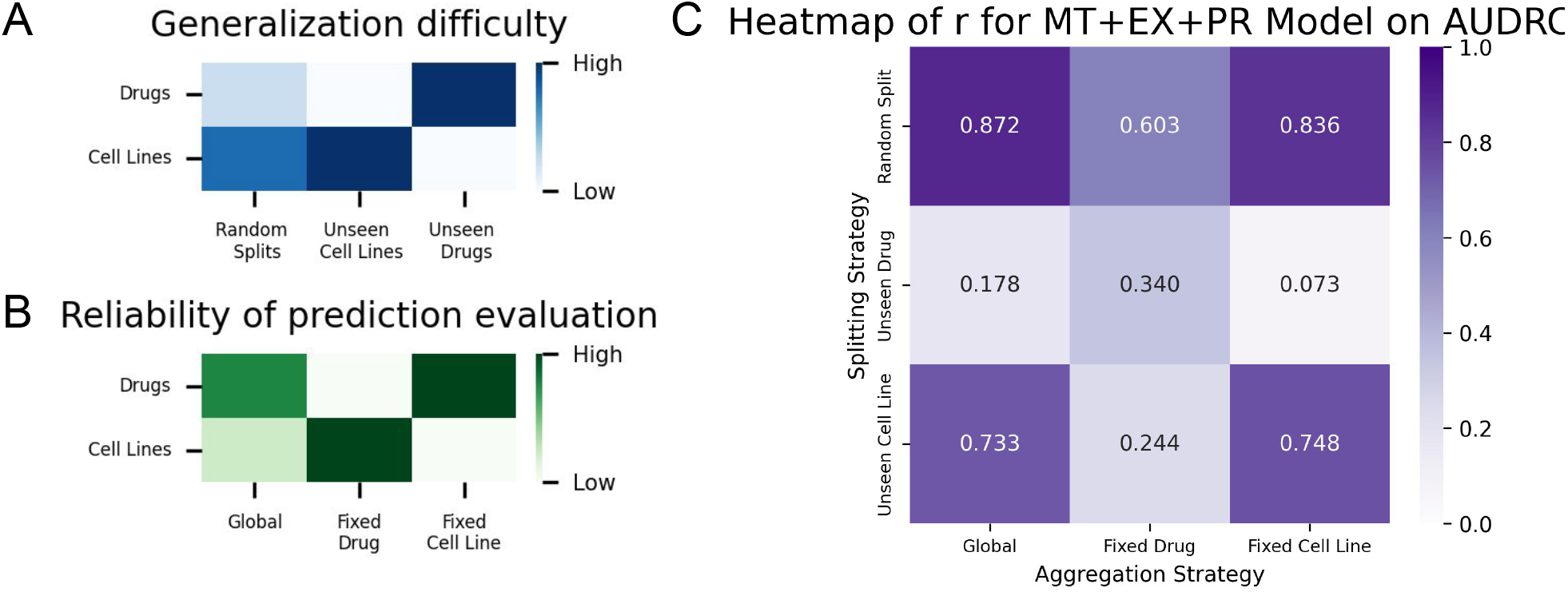
A) Heatmap *qualitatively* showing the difficulty of generalizing to either Drugs or Cell Lines with respect to different Splitting Strategies. B) Heatmap qualitatively showing how various Aggregation Strategies underscore the level of generalization achieved for different entities, such as drugs or cell lines. C) Heatmap *quantitatively* summarizing the Pearson *r* correlations of the predictions of *NxtDRP*_*MT* +*P R*+*EX*_ on AUDRC in relation to both Splitting Strategy and Aggregation Strategy.

### 2.0.2 The prediction Aggregation Strategies are crucial to evaluate different aspects of the predictions

Once the predictions are computed with one of the train/test splitting strategies described above, some performance metrics must be computed on the predictions to evaluate their level of agreement with the *ground truth* used as labels.

Typically, in ML model validation, metrics are applied “globally”, namely they are computed across all predicted values in the test set, and this is indeed the approach usually adopted by DRP methods. But what is precisely lost, in terms of analysing what the predictions mean, by using just this Global averaging? Here we look at the predictions from two other angles as well, computing the performance scores also across drugs and cell lines separately, obtaining three prediction Aggregation Strategies:

1. *Global:* This is the most common approach. The performance metrics are calculated over the entire test-set (i.e., overall correlations).
2. *Fixed-Drug:* In this Aggregation Strategy, the performance metrics are computed individually for each drug, and the resulting Fixed-Drug performance scores are then averaged over the entire test-set (across all the drugs). This Aggregation Strategy enables us to analyze the prediction quality for individual drugs independently, thereby highlighting the model’s ability to discern between the potentially unique behaviours of different cell lines.
3. *Fixed-Cell Line:* In the third Aggregation Strategy, the metrics are calculated individually for each cell line in the test set. The resulting Fixed-Cell Line performance scores are then averaged over the entire test set. This approach allows us to analyze the performance on each cell line independently from the drugs used, emphasizing the model’s ability to distinguish between different drugs when a fixed cell line is considered.

In the scientific literature, the Global aggregation is prevalent. However relying solely on this Aggregation Strategy may result in unreliable (i.e. inflated) prediction performance, depending on the dataset characteristics. The role that these Aggregation Strategies play in terms of what they precisely measure is also tightly intertwined with the Splitting Strategy being used. For a mathematical definition of these strategies, see Suppl. Section S2 Fig. 3B visually illustrates how dependable these aggregation strategies are when it comes to evaluate the generalization ability of DRP models on drugs and cell lines. Throughout the rest of the paper, we delve into this phenomenon in detail.

### 2.1 A novel non-linear data fusion method for the multi-omics prediction of drug response on cancer cell lines

To be able to run the *in-silico* experiments we needed to showcase the relevance of the various combinations of train-test splits and prediction aggregation strategies for the validation of DRP methods on cancer cell lines, we developed a novel multi-omics prediction method, called NxtDRP. It is based on the NXTfusion non-linear data fusion library proposed in [31], and allows total flexibility in testing the relevance of different types of omics data, making it particularly suitable for our analyses. The NXTfusion library generalizes the classical Matrix Factorization approach to perform inference over heterogeneous sources of information represented as Entity-Relation (ER) graphs. Each Entity in the ER graph corresponds to a class of objects (i.e. Cell Lines and Drugs), and it is internally represented by a set of latent variables that are optimized to accurately predict the target labels (i.e. IC50 values). Each -omic data matrix is added to the ER graph as a Relation connecting two Entities. For example, the Proteomics and RNA-Seq data are respectively represented as relations between the Cell Lines and the Proteins and between the Cell Lines and the Genes entities (see Fig. 6 and Methods Section 4.4 for more details).

To benchmark NxtDRP, we used the GDSC dataset, which is the most commonly used database for this task [32], due to its size and the availability of various omics to characterize the cell lines.

As mentioned before, this dataset exhibits greater variability between drugs than between cell lines. This peculiarity is shared also by the other major DRP datasets, such as CCLE and CTRP (see Fig. 4).

**Figure 4:**
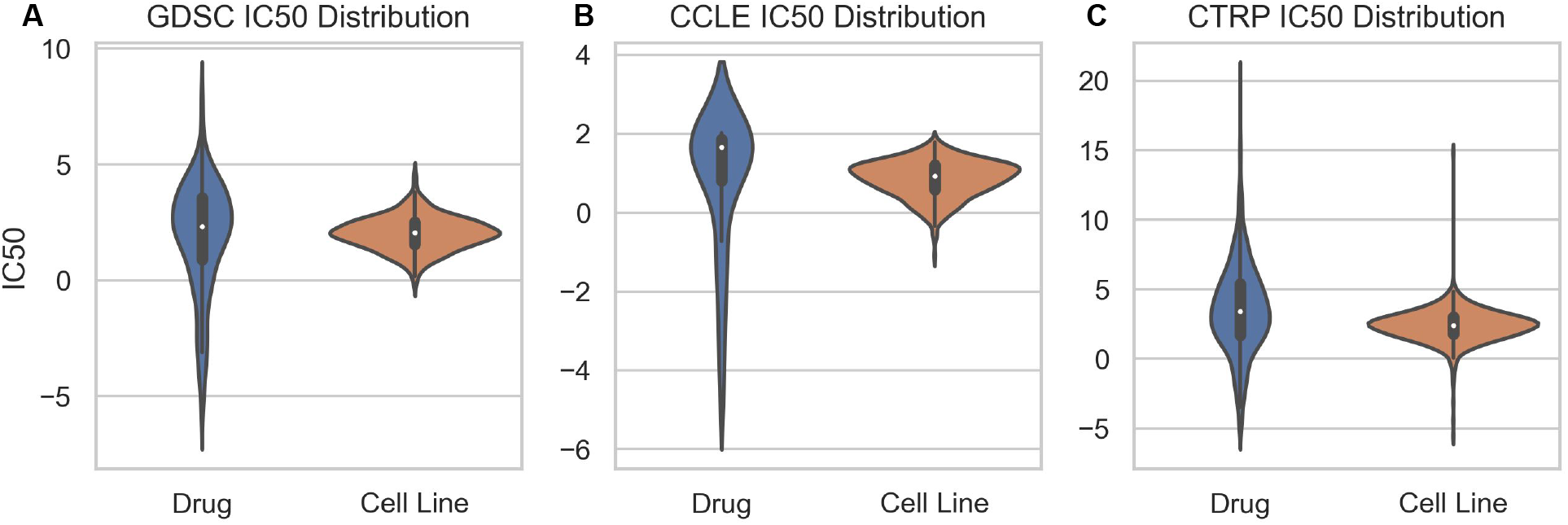
Distribution of drugs and cell lines mean IC50 values across the main DRP datasets: A) GDSC: Genomics of Drug Sensitivity in Cancer. B) CCLE: Cancer Cell Line Encyclopedia. C) CTRP: Cancer Therapeutics Response Portal.

We extracted 948 cell lines and 223 drugs, totaling 172,114 drug-response values in the form of IC50. We followed the same pre-processing steps proposed in [15] (See Methods Section 4.1 for more details). Fig. 1A, shows an overview of NxtDRP and the multi-omics data it integrates, such as RNA-Seq expression and Proteomics profiles from GDSC.

We compared the performance of NxtDRP with different ER graphs, to evaluate how the integration of different omices contribute to the prediction. The most basic ER graph involves only the Main Task (MT), namely the matrix containing the IC50 values corresponding to the Cell Line-Drug pairs available in GDSC (Fig. 6B). We then added to this minimal ER graph the available omices, one at a time, including Proteomics (PR) and RNA-Seq (EX) matrices as additional relations (see Fig. 6C-E), in an attempt to better characterize the cell lines. Throughout the paper, we refer to these relations respectively as MT, PR and EX, to indicate which are included in each model.

To make it more suitable for the DRP, we also extended the original NXTfusion library [31] by adding a Graph NN to incorporate the molecular details of the target drugs. See the Methods Section 4.3 and 4.4 for more details.

We benchmarked NxtDRP with two previously published DRP methods: tCNN [15] and GraphDRP [14]. To ensure a fair comparison, we adopted the same iterated training-test design they used, as described in the tCNN paper [15]. This means that we trained and tested NxtDRP 40 times, each time with a new selection of the train and test sets, accordingly to the chosen train-test Splitting Strategy. In each split, 90% of the dataset is selected on the basis of the Splitting Strategy adopted and is used as training set, and the remaining 10% is used as test set.

To measure the performance of the methods we adopted the Root Mean Squared Error (RMSE) and Pearson correlation metrics (*r*), averaging them over the train-test iterations (see Suppl. Mat. Section S1 for the details). More details on the validation are available in Methods.

### 2.2 The prediction of randomly selected Cell Line-Drug pairs is achieved with high accuracy

The Random Splits validation strategy measures how good a predictor is at *filling the gaps* in untested drugs-cell lines pairs. Practically, this corresponds to filling an incomplete screening on a panel of otherwise known cell lines and drugs. When validating a model following a Random Splits strategy, developers and users must know that the model is not actually tested on its ability to generalize to cell lines or drugs for which we completely lack drug-response measurements.

The results in Fig. 2A are obtained with the Random Splits validation strategy. The Global performance column indicates that NxtDRP, tCNN[15] and GraphDRP[14] are able to achieve high scores, with a Pearson correlation up to *r* = 0.93 over the entire dataset. The main reason why this setting does not present a substantial challenge is that the same drugs and cell lines can be present in both the training and test set (just not the same cell lines and drug pairs).

In Fig. 2A there is no substantial difference in performance, across all the aggregation strategies, between the model that incorporates omices such as the proteomics profile and the one that does not (see the NxtDRP_*MT*_ and NxtDRP_*MT* +*P R*+*EX*_ models). This can be explained by the fact that, in terms of information content, all the IC50 values the model observes for a given cell line and drug are sufficient to saturate the information it can learn, and therefore the additional omics data have no added value. In the Random Splits strategy, the contextual information provided by multi-omics measurements does not aid characterizing the involved entities (Drugs and Cell Lines) more than what the drug-response data alone (NxtDRP_*MT*_) already achieves.

If we focus on the other columns of Fig. 2A, we see that the Fixed-Drug performances are lower than Global performances. One probable cause of this behavior is that the model is trained *globally*, and, by minimizing its loss function, it therefore tends to capture primarily the main source of variance present in the data. As shown in Fig. 4A, in GDSC (as it is in other DRP datasets), this variance is indeed primarily driven by drugs. This is a possible reason for the poorer modeling of the cell line variability highlighted by the Fixed-Drug performance. At the same time, we can observe that Global performance and Fixed-Cell Line performance are comparable. This is explained by the fact that, while the Global performance reflects the variance both within Drugs and Cell Lines, and the Fixed-Cell Line considers only the variance within drugs, since the primary source of variance in the data is *globally* due to the drugs anyway, the two measurements are indeed extremely similar.

### 2.3 The validation on Unseen Cell Lines highlights the necessity of considering different Prediction Aggregation strategies

What happens instead if we populate the test sets used during validation only with samples coming from cell lines that are *not* present in the corresponding training sets? In real-life scenarios, this would evaluate the ability of our models to use the multi-omics data to generalize to unseen cell lines, without having observed any drug-response values on them.

#### 2.3.1 Why are Global performances of Unseen Cell Lines high even without multi-omics data?

The most noticeable aspect in Fig. 2B is that the Global performance of NxtDRP_*MT*_, which does not include any multi-omics information able to contextualize the cell lines, is already able to reach state-of-the-art performance (*r* = 0.85).

To study this perplexing behaviour we looked at the distribution of the drugs IC50 values over the cell lines. As mentioned in Section 2.2, in Fig. 4A, we see that the variability among drugs (*σ*^2^ = 2.38) is greater than that among cell lines (*σ*^2^ = 0.72). This means that, on GDSC, the predictors can already accurately model much of the variability in the data by just observing the drug responses available in the training set, regardless of the cell lines involved. This means that, if we only consider the Global aggregation of the predictions, like most of the state-of-the-art predictors do[14, 13, 15, 33, 34, 35, 36]., we remain completely unaware of the model ability to truly generalize over cell lines, which was supposed to be the goal of this Splitting Strategy.

The fact that the high Global performance in Fig. 2B are misleading becomes evident when we compare the Global and Fixed-Drug performance of NxtDRP_*MT*_. The Pearson correlation suddenly drops from *r* = 0.85 to *r* = 0, as it is supposed to be, since NxtDRP_*MT*_ with the Unseen Cell Lines Splitting Strategy *could not learn anything* about the cell lines drug-response in the test set. This drop dramatically highlights how the Global Aggregation of prediction can lead to inflated and therefore meaningless performance scores: we see a Global Pearson correlation of 0.85 despite the *actual* model complete inability to discriminate unseen cell lines (NxtDRP_*MT*_ Fixed-Drug *r* = 0).

When additional relations are added to characterise the cell lines (NxtDRP_*MT* +*EX*+*P R*_) the model reaches a Global *r* = 0.87, outperforming tCNN and GraphDRP respectively by 3.44% and 2.71% (see Suppl. Table S2).

In the Fixed-Drug settings of Fig. 2B we see that adding additional relations to the model (NxtDRP_*MT* +*EX*+*P R*_) becomes highly relevant for the performance, providing an increase in Pearson correlation, from 0.00 to 0.33, while the multi-omics contribution was only a 2% *r* increase in the Global aggregation.

#### 2.3.2 A dummy model to highlight the inadequacy of the Global performance aggregation

To make the analysis of this behavior clearer, we introduce the DummyDrugAvg model, which predicts the IC50 for a drug *d* as the *average IC50* of *d* as observed on the training set. (See Methods Section 4.5 for more details). DummyDrugAvg extremizes the situation already observed with NxtDRP_*MT*_, since by construction it cannot model or recognize different cell lines. Nonetheless, it achieves a Global Pearson correlation of 0.85 (see Fig. 2B), which is in line with all the *real* DRP methods benchmarked in the same settings. This value corresponds indeed to the correlation we measured in Fig. 1E. Similarly to NxtDRP_*MT*_, DummyDrugAvg suddenly drops to 0.0 correlation when the Aggregation Strategy switches to Fixed-Drug since it cannot predict how the same drug performs on different cell lines, but just the Global drugs trends over the entire test-set.

On the other hand, DummyDrugAvg and all the NxtDRP variants reach a high correlation (*r ≃* 0.88) in the Fixed-Cell Line aggregation performance, since the Global ranking of the drugs effectiveness is mostly conserved also within each individual cell line (see Suppl. Fig. S3). This means that generally strong drugs will be stronger than generally weaker drugs, with an average correlation across cell lines of *r* = 0.885.

### 2.4 Exploring the pitfalls of the validation on Unseen Drugs

Mirroring the previous Splitting Strategy on Unseen Cell Lines, here we predict the IC50 values for drugs that are entirely absent from the training set. To perform well in the Unseen Drugs Splitting Strategy, a DRP model should be able to generalize to drugs never seen before, relying only on biologically relevant information such as its structure. In the case of NxtDRP, the drug structure is extracted from PubChem [37] and fed to the model via a Graph NN (see Methods Section 4.4 for more details).

The accurate prediction of IC50 values for the Unseen Drugs Splitting Strategy is currently a challenge, as shown by the lack of reliable *in silico* methods for this type of validation[13, 15]. The difficulty of this task is indeed confirmed by the generally low prediction performance shown in Fig. 2C. Moreover, the variability is very high, making it impossible to directly compare models. This results from low-accuracy predictions and from the high variability of the drugs’ IC50 values, which leads to very heterogeneous subsets in each split.

The difficulty of the problem posed by this Splitting Strategy is also evident from the other prediction Aggregation Strategies shown in Fig. 2B. In the Fixed-Cell Lines aggregation, which evaluates the models’ ability to predict the dynamics between different drugs acting on the same cell line, the performance of NxtDRP_*MT*_ has a 22% drop (*r* = 0.28) with respect to Global performance. Consistently with this trend, the RMSE values in Suppl. Table S3 are generally higher than those in Suppl. Table S1 and S2.

In Fig. 2C, the Global performance is relatively similar to the Fixed-Cell Line performance. As already mentioned in Section 2.2, this is because the former, due to higher variability among drugs (see Fig. 4), tends to measure mostly how well the model approximates the dynamics of drug-related IC50 values, which is the same that the Fixed-Cell Line Aggregation Strategy *exclusively* does.

To better understand these results, we compared it with two baseline *dummy* predictors. The first is DummyCellAvg, which simply computes the mean of the IC50 values associated to each cell line *c* on the training set, and uses this mean as prediction value for any drug applied on *c* in the test set. This method achieves a Global correlation of *r* = 0.26. This value is substantially lower compared to the DummyDrugAvg performance on the unseen cell lines, which was *r* = 0.85 (see Suppl. Table S2). This is due to the fact that they both just exploit the variance in the dataset (respectively among cell lines and drugs), and the second is substantially higher than the first, as shown from the comparison of Fig. 1E and F. These plots show that the Global correlation achieved by DummyCellAvg corresponds exactly to the correlation in Fig. 1F, showcasing what a Global Aggregation Strategy truly measures when it comes to quantifying the prediction performance of DRP methods on datasets presenting a substantial structure in the data.

In the previous Section 2.3, we showed that to properly assess the models’ ability to predict previously Unseen Cell Lines, the Fixed-Drug Aggregation Strategy is the most meaningful. Analogously, here we see that to reliably evaluate the predictions on Unseen Drugs, we should use the Fixed-Cell Lines Aggregation Strategy, since it explicitly measures the ability of the model to characterise the effect of different drugs in the same condition (cell line). Indeed we see from Fig. 2C that the DummyCellAvg baseline achieves zero correlation in this Aggregation Strategy, while NxtDRP_*MT*_ with just the drug information fed through the GNN reaches *r* = 0.283.

Conversely, the Fixed-Drug performance of DummyCellAvg and NxtDRP_*MT*_ in Fig. 2C is higher to both Global and Fixed-Cell Line performance (*r ≃* 0.50). The reason for this is that the IC50 values of cell lines are similarly distributed among different drugs (*r* = 0.51, see Suppl Fig. S3). Analogously to the evidence in Section 2.3.2, this confirms that certain cell lines tend to be more sensitive to drugs while others are generally less sensitive, albeit this difference is less pronounced than the analogous effect among drugs.

The addition of omics data in the Unseen Drugs splitting provides little to no noticeable improvement in terms of performance. Indeed all the omices characterise the cell lines, and the information content is already saturated by the dose-response data.

### 2.5 In Unseen Drugs settings, the Maximum Concentration tested alone is a better predictor than ML models

The drastic drop in performance from Fig. 2A and C confirms the general observation [38, 8, 15, 14] that DRP predictors struggle to generalize to unseen drugs [13, 15], with high errors on the IC50 predictions.

Part of this issue might be related to the fact that different drugs are tested at different concentration ranges, resulting in IC50 values that are expressed within these ranges. This behavior is also visible in Fig. 1G, where we show that the Maximum Concentration (MC) at which the drugs are tested strongly correlates (*r* = 0.66) with the IC50.

To gauge how this might effect the models predictions, we ran two additional tests in which our models are provided with the MC at which each drug has been experimentally tested in order to infer the IC50. These concentration ranges are typically selected *a priori* based on existing *in vitro* and clinical data associated with each drug [23].

We first added the MC as a feature in NxtDRP (see box NxtDRP_*MT* ++*P R*+*MC*_ in Fig. 2C), by concatenating it with the drug representation generated by the GNN. This value alone increases the Global Pearson correlation by 67% (*r* = 0.60). To further investigate the role of MC in relation to the other inputs fed to NxtDRP_*MT* +*MC*_, we designed an additional baseline model, called DummyMC. For each drug *d*, DummyMC predicts its IC50 by simply outputting the MC tested for *d*. This trivial approach outperforms any other DRP predictor in the Global prediction Aggregation Strategy (*r* = 0.61) without even requiring any type of modeling of the cell line and the drug.

If we look at the Fixed-Drug performance in Fig. 2C, we see that the DummyMC Pearson correlation drops to almost zero, since it always outputs the same IC50 value to each drug *d*, regardless of the cell line involved. Conversely, NxtDRP_*MT* +*P R*+*MC*_ achieves *r* = 0.56.

DummyMC simply reflects the assumptions on which the *in vitro* experiments that DRP models wish to emulate, and it could therefore considered a *baseline* for benchmarking DRP performance. Unfortunately, as we show in Fig. 2C, current DRP methods are not able to surpass this baseline.

## 3 Discussion

In this paper we highlight two issues that, to the best of our knowledge, have not been adequately addressed in the DRP field. We believe that the development of increasingly sophisticated DRP models cannot ignore the need for a validation protocol that truly assesses their generalization capability. The critical points we have addressed concern the use of IC50 as the target label and the standard approach adopted for the validation of DRP models.

### 3.1 Adopting the Area Under the Dose-Response Curve as prediction label to overcome the IC50 Limitations

As highlighted in Section 2.5, using IC50 as the target for prediction creates situations in which much of the variance in the data is due to the MC used to test different drugs, while the *truly* interesting variance due to the actual biological diversity of the cancer cell lines targeted by the drugs is negligible in comparison. We showed how IC50 values strongly correlate with MC (see Fig. 1G), and how this becomes problematic since a baseline represented by the mere MC value (DummyMC) results in higher performance than advanced DRP models. What makes this correlation even more problematic is the fact that it is subtly influenced by the assumptions underlying the IC50. IC50 is based on the existence of a concentration inhibiting 50% of cells [39]. In reality, 62% of the IC50 values in GDSC are higher than the actual MC tested for the target drug, meaning that most of the IC50 values *have never been actually experimentally observed, and it is interpolated from the dose-response sigmoid instead*. This is even more striking on the CCLE dataset, since when the IC50 exceeded the MC tested (the 55% of the cases), the IC50 was approximated *by using the MC value instead*.

To overcome these IC50-related issues, we could try to adopt an alternative prediction label to be used to train DRP methods. Ideally, such a target label should represent both the drugs potency, represented by metrics such as IC50, and efficacy, which denotes the maximum effect or response that the drug can produce [40, 38].

An option in this sense is to use the Area Under the Dose-Response Curve (AUDRC) (see Fig. 1B). While it is commonly referred to as AUC in the chemistry literature, we refer to it as AUDRC in this paper to avoid confusion with the Area Under the Receiver Operating Characteristic curve (AUC, or AUROC), which commonly used in ML [39]. The AUDRC has already been proposed as target label for DRP methods [39, 41]: an AUDRC of 1 means that the drug is not able to inhibit the viability of cancer cell, even at the MC tested, while an AUDRC close to zero indicates that the drug is extremely effective against cancer cells, even at the lowest concentrations tested.

The AUDRC therefore effectively *detaches* the *concentration ranges* tested for each drug to the notion of their effectiveness as anticancer drug, since these ranges now determine the limits of the calculated area, as shown in Fig. 1B. The AUDRC is essentially a comprehensive summary of the dose-response curve based on the tested concentrations, and it relies unambiguously on the assumptions made in the *in vitro* experiments, so we endorse the use of AUDRC to avoid the subtle and potentially misleading dependencies between IC50 and MC that we have shown in this paper.

### 3.2 A more robust validation protocol for DRP methods based on AUDRC and aggregation strategies

When using the AUDRC, there is still a difference in variability between drugs and cell lines, albeit to a lesser extent (see Fig. 5A), since the AUDRC values are more dependent on the drugs than on the cell lines. For this reason, changing the DRP target label from IC50 to AUDRC alone is not sufficient for a completely reliable validation. It is also necessary to couple it with the proper prediction Aggregation Strategies, to avoid the issues highlighted in Sections 2.3 and 2.4, since also the AUDRC Global performance could be biased by the fact that it mostly describes the model’s ability to explain variance among drugs rather than among cell lines.

**Figure 5:**
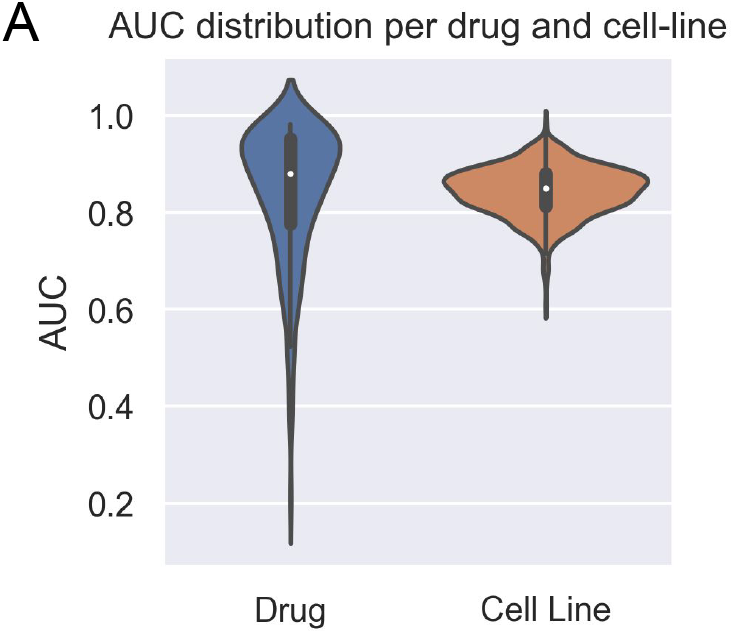
A) Mean AUDRC values distribution of drugs and cell lines.

**Figure 6:**
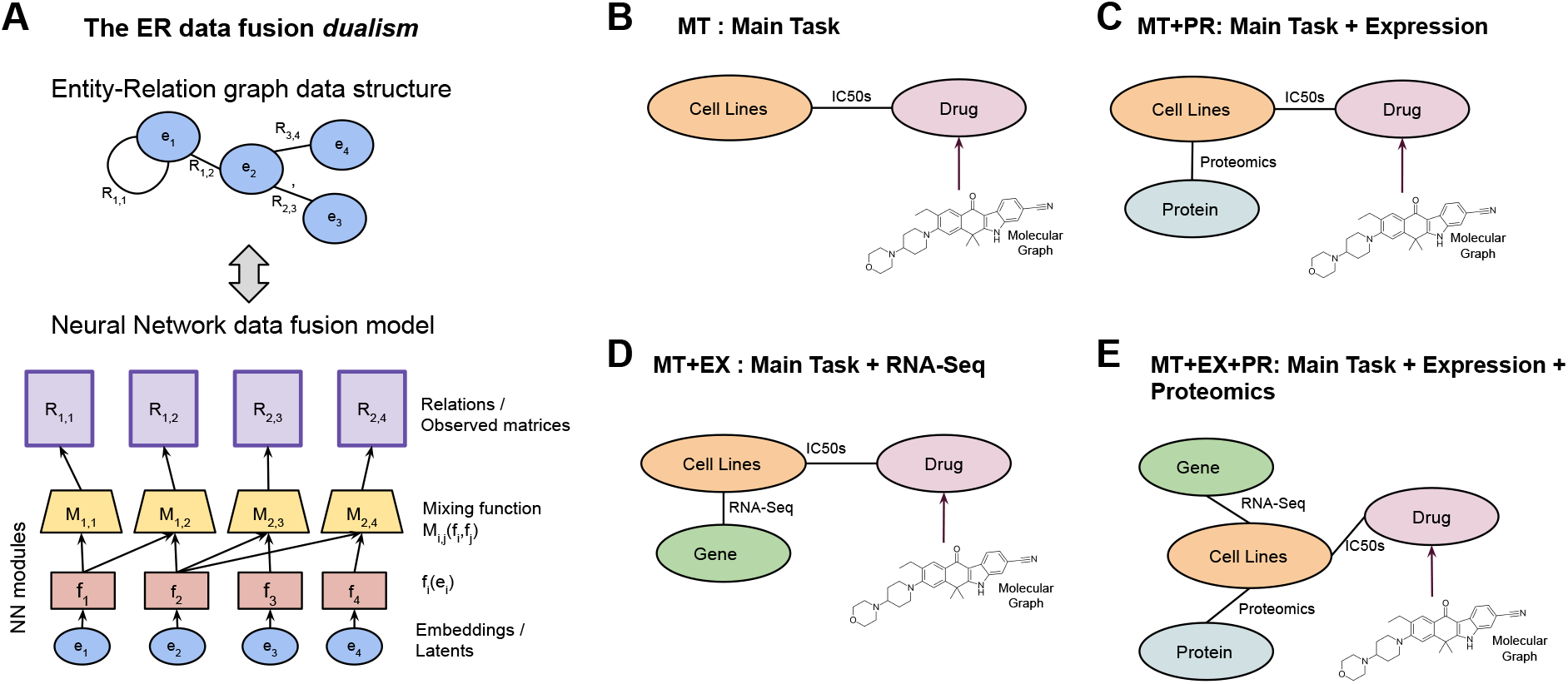
A) Representation of the dualism of the abstract representation of the data in an Entity-Relation graph and the NXTFusion schematization of the non-linear inference over that ER graph. B) ER Graph representing only the Main Task that is the DRP on IC50 values. C) ER Graph representing the Main Task with the addition of RNA-Seq expression relation. D) ER Graph representing the Main Task with the addition of Proteomics relation. E) ER graph representing the Main Task with both Proteomics and RNA-Seq relations.

To provide a complete solution to these issues, here we propose a novel *validation protocol* for DRP methods, which is free from the issues we highlighted so far:

1. Use AUDRC as prediction label, instead of the IC50.
2. Aggregate the performance using the Fixed-Drug and Fixed-Cell Lines strategies (see Section 2.0.2). In particular, the Fixed-Drug strategy *describes the model ability to discriminate between cell lines*, and vice versa, the Fixed-Cell Line performance *indicates how well the model correctly distinguishes between different drugs*.
3. In the case of a validation with Unseen Drugs train-test splits, the most relevant Aggregation Strategy will be Fixed-Cell Line. Conversely, to evaluate the generalization ability in the Unseen Cell Lines splits, the main prediction Aggregation Strategy should be Fixed-Drug.

#### 3.2.1 A reproducible and fair benchmark for DRP methods

To showcase the novel validation procedure we propose, we tested NxtDRP in these settings. The results are shown in Supplementary Material Section S4. In the Random Splits setting, (see Suppl. Table S4), the performances are similar to the ones obtained with IC50. The raw predictions are available at github.com/codicef/DRPValidation, to allow future DRP methods developed on GDSC data to benchmark their results with NxtDRP and our validation protocol.

The Unseen Cell Lines train-tests splits (see Suppl. Table S6) achieve a Global correlation of *r* = 0.733, which is indeed biased by the prominence of the drug variability. Similarly to the IC50 settings in Suppl. Table S2, the Fixed-Drug performance show a drastic drop in performance (*r* = 0.244), indicating an actually lower ability to truly predict the variability of cell lines.

In the Unseen Drug train-test splits (see Suppl. Table S5), NxtDRP_*MT*_ struggles to generalize, obtaining Global correlation of *r* = 0.171 and a Fixed-Cell Lines correlation of *r* = 0.081. This is in line with the IC50-based performance we showed in Suppl. Table S3. It indicates that generalizing to never seen before drugs is currently an insurmountable challenge for DRP methods, even if the actual drug molecular structure is provided as input. This might be due to the fact that the currently available sample size does not allow the model to learn any useful relation between the drug molecular structures and the biology of the cancer cells, since they both are immense spaces.

To see whether a mitigation for this problem could be already in sight, we tried to pre-train the GNN in an unsupervised manner with 20,000 drug structures from NCI60[3], using this as initialization for the actual training of NxtDRP on GDSC data. Unfortunately this did not produce higher results as even on the NCI60 dataset alone, we encountered challenges in achieving satisfactory levels of performance. Further refinements and adjustments may be necessary to enhance the effectiveness of the pre-training approach and subsequent model training.

### 3.3 The long overlooked role of the Aggregation Strategies

Casting aside the discourse related to the target label of choice for a moment, our study shows that the validation of DRP methods is hindered by the fact that the *only* Aggregation Strategy currently used in literature (Global) biases the performance towards the entity showing greater variance, which happen to be the drugs entity in GDSC and CCLE. This leads to models that reach completely misleading high Global performance, even though they are actually unable to generalize on these entities.

The differentiation of the Aggregation Strategies that we propose addresses this issue. In addition to the Global aggregation, Fixed-Drug and Fixed-Cell Line aggregations must be used to assess the true performance of DRP methods. They assess the ability of DRP models to generalize to unseen cell lines (Fixed Drug Aggregation) and to unseen drugs (Fixed Cell Line Aggregation), as we showed in Sections 2.3 and 2.4.

Aggregation Strategies are also relevant to evaluate the predictions in the Random Splits setting, as they provide an overview of the degree of generalization achieved independently for Drugs and Cell Lines.

## 4 Methods

### 4.1 Datasets

In this study we used as dataset containing cancer cell lines drug sensitivity measurements: GDSC v. 6.0 [5]. It contains 1,074 cell lines and 224,510 IC50 measurements of 265 drugs. For each cell line, the following omices are available: whole genome sequencing, transcriptomics, proteomics, copy number variation and methylation data. Drugs are identified by their PubChem IDs, facilitating the retrieval of their chemical structures.

GDSC dataset contain drug dose-response data measurements. They are obtained using fluorescence signal intensities, testing 9 different concentrations per drug with 2-fold dilution series [5]. The dose-response curve is then fitted by a nonlinear mixed effect model on a sigmoid curve. The IC50 values we use as prediction labels are therefore the results of the interpolation of the dose-response curve with the sigmoid curve (see Fig 5A). These IC50 values are therefore subject to noise, due for example to high experimental variability that may cause a poor fitting, which is estimated by an RMSE value. We excluded IC50 values with an RMSE *>* 0.3 from the analysis.

In order to be able to fairly compare our results with the already existing approaches, we adopted the preprocessing used in [15]. First, we considered only compounds with a PubChem ID, obtaining a final dataset of 223 drugs and 948 cell lines, with a total of 172114 drug-response IC50 values. Second, we rescaled them between 0 and 1 in the following way 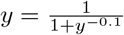 [15].

### 4.2 Representing drug molecular structures in a Machine Learning-understandable format

Each drug in GDSC is uniquely identifyied by a Pubchem ID. We retrieved their Simplified Molecular Input Line Entry System (SMILES) representation from Pubchem [37]. However, this representation cannot be meaningfully used as input in a DL model as it is. We therefore transformed them into graphs that represent the drugs’ molecular structures, using PyTorch Geometric (PyG) [42] to feed them as input to a Graph NN.

Each drug is represented by a graph in which nodes correspond to an atom, and it is described by the following set of features: atomic symbol (one-hot encoding), atomic number, atomic degree, atomic formal charge, atom in a ring, atom radical electrons, atom hybridization state, and atom aromaticity. Each edge in this graph correspond to a chemical bond between atoms.

### 4.3 An Entity-Relation Data fusion model to predict the cancer cell lines drug sensitivity

Predicting the response of cancer cell lines to anticancer drugs requires integrating heterogeneous sources of information, such as the omices available for each cell line and the molecular information related to each drug. To build a model able to do that, we started from our data fusion framework NXTFusion [31], which allows us to describe these heterogeneous data sources within an Entity-Relation (ER) Graph (see Fig. 6A), which can be intuitively thought as a relational database on which it is possible to perform inference.

An ER graph consist of two key elements: a set of entities denoted as *E*, which represent classes of objects, and a set of relations *R* that specify how *pairs* of entities are interconnected. Each entity has a specific cardinality that corresponds to the number of instances present in the data available. For example, the Cell Line entity on GDSC data has 948 instances.

From a modeling perspective, the entities are represented by a set of trainable latent variables *e*_*i*_. Each observed data matrix *R*_*ij*_ (i.e. proteomics data) correspond to a relation between a pair of entities (*i, j*).

The inference is performed globally over the ERG by finding the embeddings *e*_*i*_ and *e*_*j*_ that minimize the reconstruction error on each observed matrix (relation) *R*_*ij*_. The loss function to be minimized for each *R*_*ij*_ is therefore *L*_*ij*_(*R*_*ij*_, *M*_*ij*_(*f*_*i*_(*e*_*i*_), *f*_*j*_*e*_*j*_)) where *L*_*ij*_ is the relation-specific loss function, *f*_*i*_ is an entity specific differentiable function (such as a feed-forward NN), and *M*_*ij*_(., .) a relation-specific mixing function (such as bilinear layer) [31]. The loss function *L*_*ij*_ can match the type of data contained into each relation *R*_*ij*_, depending on whether it is a classification, regression or multi-class prediction task (see Fig. 6).

Since the ER graph might contain an arbitrary number of relations between entities, the Global objective function to be optimized is the following

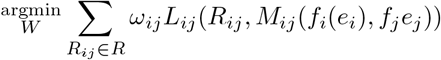

where *W* is the set of trainable parameters of each NN module in *f* and *M* functions. *ω* is a scale factor meant to ensure that different relations have a comparable weight in the Global loss (see [31] for more details).

Essentially, NXTfusion achieves data fusion through learning several tasks concurrently. The auxiliary tasks help the inference of the main task by providing additional context, which acts as an *informed regularization* [31], therefore helping the generalization on the main task by ensuring the convergence to more informative latent representations of the entities involved in the ER graph [27].

### 4.4 Entity-Relation Graph Inference for DRP

In the context of the GDSC data, we devised relations between the cell lines, drugs, proteins and genes entities (see Fig. 1A). The number of instances in each entity depends on the available data points in the datasets.

The Cell Line-Drug relation is based on the drug-response values (IC50) from GDSC. It contains 172114 IC50 values between 948 cell lines and 223 drugs. The Cell Line/Protein relation is based on quantitative proteomics intensity values. We scaled the data to be in the [0,1] range. This relation contains 4538041 data points between 874 cell lines and 8457 proteins. The relation between Cell Lines and Genes is based on RNA-Seq data and it contains 20080264 transcript per million (TPM) measurements. We scaled these values to be in the [0,1] range. This relation contains involves 912 Cell Lines and 36447 genes.

The last crucial piece in the picture is the contextualization of the drugs. The detailed molecular description is available through their PubChem structures. We therefore decided not to use a latent representation (embedding) to represent them, but to extend the concept of *side information* (i.e. features in the Matrix Factorization jargon [31, 43]) to allow this information to be introduced in to our ER Graph in an *end-to-end* fashion with a Graph NN directly modeling their molecular structures.

We retrieved the molecular graph of each drug from PubChem [37] (see Methods Section 4.2) and we processed them with a Graph NN whose output is directly connected to the Cell Line-Drug relation (see Fig. 6B-E). The Graph NN consists of 4 Graph Attention Layers (GAT) (v. 2 [44]), followed by a final Global sum pooling step. The final pooling is necessary to provide a final latent representation describing each drug that is independent of each drug’s actual number of atoms.

The method along with the code for its implementation is available at github.com/codicef/NxtDRP to reproduce the results.

### 4.5 Dummy models

The dummy models in this paper are categorized into two groups. The first group is based on simple transformations of the IC50 values from the dataset, while the second utilizes external information, such as the maximum concentration (MC) at which the drug was tested.

For the first category, these models employ IC50 data based on the specific validation performed. For Unseen Cell Lines, the average IC50 values for each drug (DummyDrugAvg) are used, excluding data from cell lines in the test set. Similarly, for Unseen Drugs, the average IC50 values for cell lines (DummyCellAvg) are utilized, excluding data from drugs in the test set. In the Random Splits strategy, a basic linear regression model (DummyLR) trained on concatenated one-hot identifiers for drugs and cell lines.

As for the dummy model based on MC, it simply uses the MC value at which each drug was tested (DummyMC). This information is known a priori from the screening experiments that the DRP models aim to replicate.

## Acknowledgements

FC thanks E. Salvadori for her support and constructive discussions. FC is also grateful to T. Sanavia and G. Birolo for their help and scientific insights during the project. Special thanks to S. Zanella for assistance with the graphics. DR is grateful to A. L. Mascagni.

