## Supplementary Material for "The Specification Game: Rethinking the Evaluation of Drug Response Prediction for Precision Oncology"

July 13, 2024

#### S1 Evaluation Metrics

The RMSE measure the difference between the actual values $y$ and the predicted ones $\hat{y}$ and it is computed as follows:

$$\text{RMSE} = \sqrt{\frac{1}{n}\sum_i (y_i - \hat{y}_i)^2}$$

The Pearson correlation is a measure of the linear correlation between two sets of values and it is defined by the following formula

$$r = \frac{cov(y, \hat{y})}{\sigma_y \sigma_{\hat{y}}}$$

#### S2 Aggregation Strategies

Given $y_{d,c}$ as the true target label for Drug $d$ and Cell Line $c$, and $\hat{y}_{d,c}$ as the corresponding predicted value.

Let $DR$ be the set containing the tuples $(d, c)$ represented in the test-set, $D = \{d|(d, c) \in DR\}$ be the set of the Drugs, and $C = \{c|(d, c) \in DR\}$ be the set of the Cell Lines.

Let $m(y, \hat{y})$ be any regression scoring function between a set of true labels $y$ and the corresponding set of predicted labels $\hat{y}$.

- *Global Aggregation*: The Global performance is calculated using all pairs $(d, c) \in DR$ in the test set:

$$m_{\text{Global}} = m(\{y_{d,c}|(d, c) \in DR\}, \{\hat{y}_{d,c}|(d, c) \in DR\})$$

- *Fixed-Drug*: The performance for each drug is computed individually, and then averaged across all drugs:

$$m_{\text{Fixed-Drug}} = \frac{1}{|D|}\sum_{d^* \in D} m(\{y_{d,c}|(d, c) \in DR, d = d^*\}, \{\hat{y}_{d,c}|(d, c) \in DR, d = d^*\})$$

- *Fixed-Cell Line*: The performance for each cell line is computed individually, and then averaged across all cell lines:

$$m_{\text{Fixed-Cell Line}} = \frac{1}{|C|} \sum_{c^* \in C} m(\{y_{d,c} | (d,c) \in DR, c = c^*\}, \{\hat{y}_{d,c} | (d,c) \in DR, c = c^*\})$$

### S3 Performance tables for IC50 as prediction label

| Entities | Global $r$ | RMSE | Fixed Drug $r$ | RMSE | Fixed Cell Line $r$ | RMSE |
|---|---|---|---|---|---|---|
| DummyLR | $0.884 \pm 0.00$ | $0.031 \pm 0.00$ | $0.501 \pm 0.01$ | $0.029 \pm 0.00$ | $0.868 \pm 0.00$ | $0.030 \pm 0.00$ |
| tCNN | $0.910 \pm 0.00$ | $0.028 \pm 0.00$ | $0.591 \pm 0.01$ | $0.026 \pm 0.00$ | $0.896 \pm 0.00$ | $0.027 \pm 0.00$ |
| GraphDRP | $0.929 \pm 0.00$ | $0.025 \pm 0.00$ | $0.687 \pm 0.01$ | $0.024 \pm 0.00$ | $0.915 \pm 0.00$ | $0.024 \pm 0.00$ |
| MT | $0.929 \pm 0.00$ | $0.025 \pm 0.00$ | $0.668 \pm 0.02$ | $0.024 \pm 0.00$ | $0.915 \pm 0.00$ | $0.024 \pm 0.00$ |
| MT+EX | $0.931 \pm 0.00$ | $0.025 \pm 0.00$ | $0.675 \pm 0.01$ | $0.024 \pm 0.00$ | $0.917 \pm 0.00$ | $0.024 \pm 0.00$ |
| MT+PR | $0.933 \pm 0.00$ | $0.025 \pm 0.00$ | $0.683 \pm 0.02$ | $0.023 \pm 0.00$ | $0.919 \pm 0.00$ | $0.023 \pm 0.00$ |
| MT+EX+PR | $0.930 \pm 0.00$ | $0.025 \pm 0.00$ | $0.672 \pm 0.01$ | $0.024 \pm 0.00$ | $0.916 \pm 0.00$ | $0.024 \pm 0.00$ |

Table S1: Table showing the benchmark of the tCNN, GraphDRP and NxtDRP variants on GDSC, with Random Splits and the three Aggregation Strategies. NxtDRP with MT, PR and EX refer to the omices used (respectively none, Proteomics and Transcriptomics, see Fig. 5 B-E for details). DummyLR is a baseline model that corresponds to a Linear Regression applied to concatenated one-hot encodings of drugs and cell lines.

| Entities | Global $r$ | RMSE | Fixed Drug $r$ | RMSE | Fixed Cell Line $r$ | RMSE |
|---|---|---|---|---|---|---|
| DummyDrugAvg | $0.849 \pm 0.01$ | $0.035 \pm 0.00$ | $0.000 \pm nan$ | $0.033 \pm nan$ | $0.877 \pm 0.01$ | $0.034 \pm 0.00$ |
| tCNN | $0.841 \pm 0.01$ | $0.036 \pm 0.00$ | $0.137 \pm 0.05$ | $0.035 \pm 0.00$ | $0.858 \pm 0.00$ | $0.036 \pm 0.00$ |
| GraphDRP | $0.847 \pm 0.01$ | $0.036 \pm 0.00$ | $0.152 \pm 0.06$ | $0.034 \pm 0.00$ | $0.856 \pm 0.01$ | $0.035 \pm 0.00$ |
| MT | $0.850 \pm 0.01$ | $0.036 \pm 0.00$ | $0.000 \pm 0.01$ | $0.034 \pm 0.00$ | $0.879 \pm 0.01$ | $0.035 \pm 0.00$ |
| MT+EX | $0.867 \pm 0.01$ | $0.034 \pm 0.00$ | $0.302 \pm 0.04$ | $0.033 \pm 0.00$ | $0.886 \pm 0.01$ | $0.033 \pm 0.00$ |
| MT+PR | $0.869 \pm 0.01$ | $0.034 \pm 0.00$ | $0.332 \pm 0.04$ | $0.032 \pm 0.00$ | $0.887 \pm 0.01$ | $0.033 \pm 0.00$ |
| MT+EX+PR | $0.870 \pm 0.01$ | $0.034 \pm 0.00$ | $0.327 \pm 0.04$ | $0.032 \pm 0.00$ | $0.888 \pm 0.01$ | $0.033 \pm 0.00$ |

Table S2: Table showing the benchmark of tCNN, GraphDRP and NxtDRP variants on GDSC, with Unseen Cell Line Splitting Strategy and with the three Aggregation Strategies. NxtDRP with MT, PR and EX refer to the omices used (respectively none, Proteomics and Transcriptomics, see Fig. 5 B-E for details). DummyDrugAvg model predicts a drug's IC50 value as the average IC50 observed for that drug in the training dataset.

| Entities | Global $r$ | RMSE | Fixed Drug $r$ | RMSE | Fixed Cell Line $r$ | RMSE |
|---|---|---|---|---|---|---|
| DummyCellAvg | $0.264 \pm 0.04$ | $0.064 \pm 0.01$ | $0.501 \pm 0.02$ | $0.057 \pm 0.01$ | $0.000 \pm nan$ | $0.046 \pm nan$ |
| Dummy(MC) | $0.611 \pm 0.14$ | $0.051 \pm 0.00$ | $0.019 \pm 0.11$ | $0.049 \pm 0.01$ | $0.640 \pm 0.19$ | $0.051 \pm 0.00$ |
| tCNN | $0.353 \pm 0.12$ | $0.064 \pm 0.01$ | $0.450 \pm 0.03$ | $0.057 \pm 0.00$ | $0.278 \pm 0.16$ | $0.063 \pm 0.01$ |
| GraphDRP | $0.339 \pm 0.20$ | $0.068 \pm 0.01$ | $0.420 \pm 0.08$ | $0.059 \pm 0.01$ | $0.278 \pm 0.25$ | $0.067 \pm 0.01$ |
| MT | $0.363 \pm 0.17$ | $0.070 \pm 0.01$ | $0.508 \pm 0.03$ | $0.060 \pm 0.01$ | $0.283 \pm 0.20$ | $0.069 \pm 0.01$ |
| MT+EX | $0.339 \pm 0.20$ | $0.068 \pm 0.01$ | $0.500 \pm 0.03$ | $0.058 \pm 0.01$ | $0.241 \pm 0.23$ | $0.067 \pm 0.01$ |
| MT+PR | $0.320 \pm 0.20$ | $0.070 \pm 0.01$ | $0.506 \pm 0.03$ | $0.060 \pm 0.01$ | $0.235 \pm 0.23$ | $0.069 \pm 0.01$ |
| MT+EX+PR | $0.319 \pm 0.20$ | $0.068 \pm 0.01$ | $0.498 \pm 0.03$ | $0.059 \pm 0.01$ | $0.217 \pm 0.24$ | $0.067 \pm 0.01$ |
| MT+MC | $0.590 \pm 0.10$ | $0.057 \pm 0.01$ | $0.495 \pm 0.02$ | $0.051 \pm 0.00$ | $0.549 \pm 0.13$ | $0.056 \pm 0.01$ |
| MT+MC+EX | $0.602 \pm 0.11$ | $0.058 \pm 0.01$ | $0.499 \pm 0.03$ | $0.051 \pm 0.00$ | $0.563 \pm 0.13$ | $0.057 \pm 0.01$ |
| MT+MC+PR | $0.602 \pm 0.14$ | $0.058 \pm 0.01$ | $0.509 \pm 0.02$ | $0.052 \pm 0.00$ | $0.560 \pm 0.18$ | $0.057 \pm 0.01$ |
| MT+MC+EX+PR | $0.568 \pm 0.14$ | $0.061 \pm 0.01$ | $0.500 \pm 0.04$ | $0.053 \pm 0.01$ | $0.528 \pm 0.17$ | $0.060 \pm 0.01$ |

Table S3: Table showing the benchmark of tCNN [1], GraphDRP [2] and NxtDRP variants on IC50 values of GDSC, with Unseen Drug Splitting Strategy and with the three Aggregation Strategies. NxtDRP with MT, PR and EX refer to the omices used (respectively none, Proteomics and Transcriptomics, see Fig. 5 B-E for details). DummyCellAvg model predicts a cell line's IC50 value as the average IC50 observed for that cell line in the training dataset. data. The DummyMC model corresponds to the Maximum Concentration tested for a given drug.

#### S3.1 Unseen Cell Line-Drug paired validation

In unseen Cell Line-Drug paired validation, the model must be able to generalize its predictions concerning not only new drugs but also new cell lines, all without having the ability to observe how they behave in terms of drug response measurements.

Our model achieves limited performance, in terms of Pearson's r, ranging from 0.26 to 0.50.

| | $r$ | RMSE |
|---|---|---|
| MT | 0.26 | 0.14 |
| MT+EX | 0.50 | 0.06 |
| MT+PR | 0.49 | 0.06 |
| MT+PR+EX | 0.50 | 0.06 |

Table S4: Performance Metrics for unseen Cell Line-Drug paired validation, the Global aggregation strategy is reported.

### S4 Performance tables for AUDRC as prediction label

| | Global | | Fixed Drug | | Fixed Cell Lines | |
|---|---|---|---|---|---|---|
| Entities | $r$ | RMSE | $r$ | RMSE | $r$ | RMSE |
| MT | 0.862(±0.00) | 0.102(±0.00) | 0.596(±0.21) | 0.087(±0.05) | 0.826(±0.18) | 0.092(±0.04) |
| MT+EX | 0.871(±0.00) | 0.099(±0.00) | 0.605(±0.21) | 0.085(±0.05) | 0.836(±0.17) | 0.089(±0.04) |
| MT+PR | 0.869(±0.00) | 0.100(±0.00) | 0.607(±0.21) | 0.085(±0.05) | 0.834(±0.17) | 0.090(±0.04) |
| MT+PR+EX | 0.872(±0.00) | 0.098(±0.00) | 0.603(±0.21) | 0.085(±0.05) | 0.836(±0.18) | 0.089(±0.04) |

Table S5: Table showing the benchmark of the tCNN, GraphDRP and NxtDRP variants on AUDRC values of GDSC, with Random Splits and the three Aggregation Strategies. NxtDRP with MT, PR and EX refer to the omices used (respectively none, Proteomics and Transcriptomics, see Fig. 5 B-E for details).

| | Global | | Fixed Drug | | Fixed Cell Lines | |
|---|---|---|---|---|---|---|
| Entities | $r$ | RMSE | $r$ | RMSE | $r$ | RMSE |
| MT | 0.171(±0.13) | 0.210(±0.03) | 0.326(±0.19) | 0.173(±0.12) | 0.081(±0.26) | 0.197(±0.07) |
| MT+EX | 0.142(±0.13) | 0.213(±0.03) | 0.329(±0.20) | 0.173(±0.12) | 0.042(±0.26) | 0.200(±0.07) |
| MT+PR | 0.139(±0.14) | 0.223(±0.03) | 0.328(±0.19) | 0.182(±0.13) | 0.050(±0.27) | 0.211(±0.08) |
| MT+PR+EX | 0.178(±0.14) | 0.210(±0.03) | 0.340(±0.20) | 0.173(±0.12) | 0.073(±0.27) | 0.197(±0.07) |

Table S6: Table showing the benchmark of the tCNN, GraphDRP and NxtDRP variants on AUDRC values of GDSC, with Unseen Drug Splitting Strategy and the three Aggregation Strategies. NxtDRP with MT, PR and EX refer to the omices used (respectively none, Proteomics and Transcriptomics, see Fig. 5 B-E for details).

| | Global | | Fixed Drug | | Fixed Cell Lines | |
|---|---|---|---|---|---|---|
| Entities | $r$ | RMSE | $r$ | RMSE | $r$ | RMSE |
| MT | 0.704(±0.02) | 0.142(±0.00) | 0.017(±0.13) | 0.120(±0.07) | 0.737(±0.13) | 0.135(±0.04) |
| MT+EX | 0.729(±0.01) | 0.138(±0.00) | 0.237(±0.18) | 0.118(±0.07) | 0.746(±0.12) | 0.133(±0.03) |
| MT+PR | 0.737(±0.01) | 0.136(±0.00) | 0.241(±0.20) | 0.117(±0.07) | 0.758(±0.12) | 0.130(±0.04) |
| MT+PR+EX | 0.733(±0.01) | 0.137(±0.00) | 0.244(±0.19) | 0.118(±0.07) | 0.748(±0.11) | 0.132(±0.03) |

Table S7: Table showing the benchmark of the tCNN, GraphDRP and NxtDRP variants on AUDRC values of GDSC, with Unseen Cell Line Splitting Strategy and the three Aggregation Strategies. NxtDRP with MT, PR and EX refer to the omices used (respectively none, Proteomics and Transcriptomics, see Fig. 5 B-E for details).

#### S5 CCLE Dataset Characteristics

A)

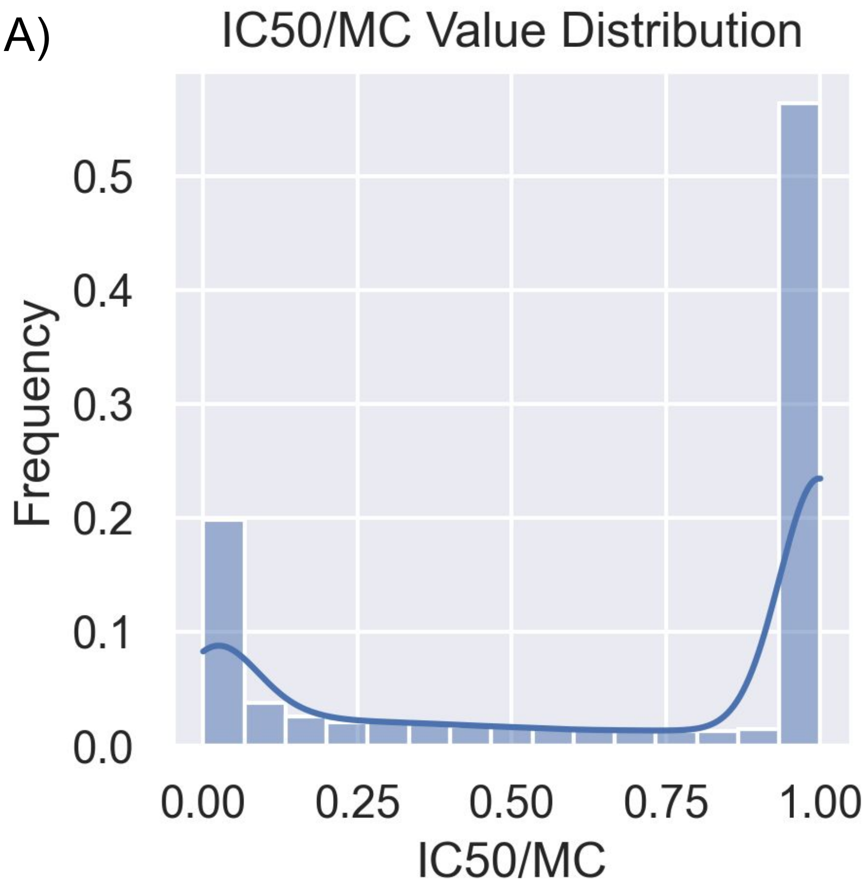


Figure S1: A) CCLE: This histogram illustrates the frequency distribution of the ratios of IC50 values/MC, with a significant concentration of higher ratios indicating that most of the IC50 values are approximated to the maximum concentration tested.

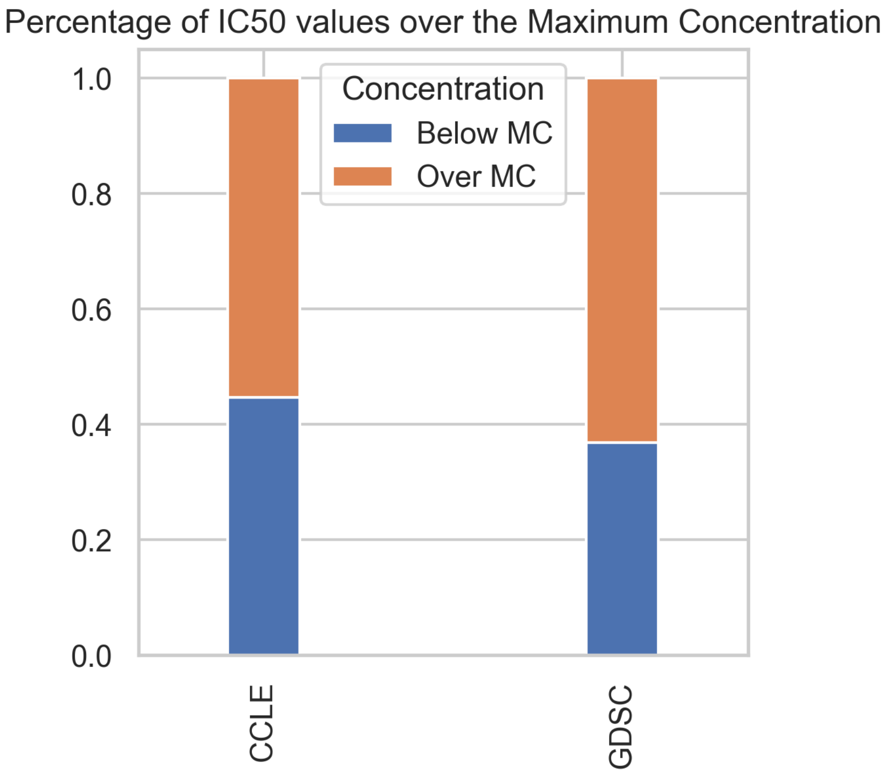


Figure S2: This bar chart shows the percentage of IC50 values relative to the maximum concentration (MC) in the CCLE and GDSC datasets, categorized into 'Below MC' and 'Over MC'.

#### S6 Dataset characteristics

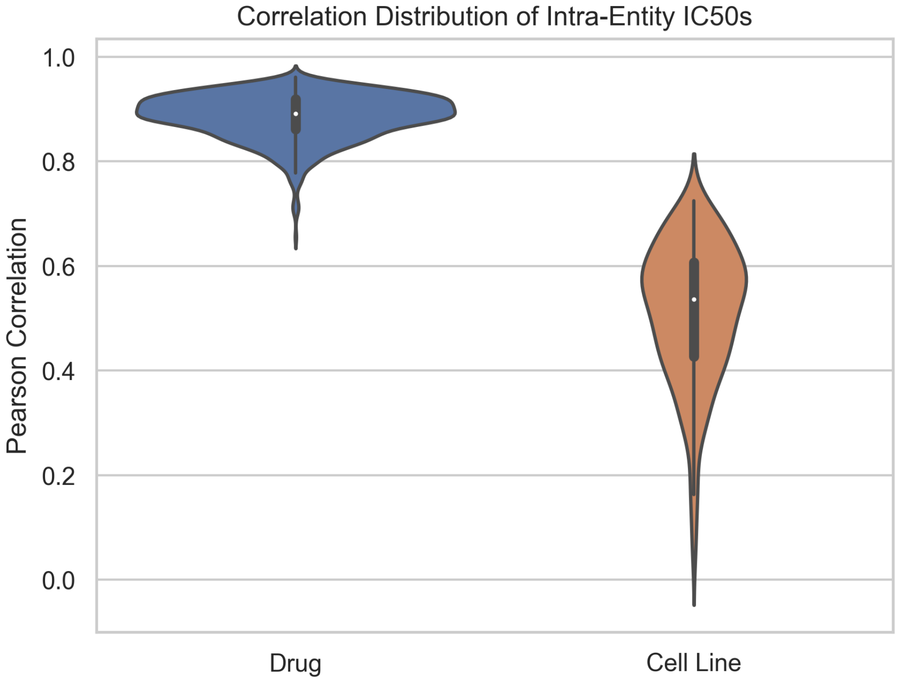


Figure S3: This violin plot illustrates the distribution of Pearson correlations between average IC50 values for each drug or cell line and the respective IC50s within each entity.
